# Spatial and phylogenetic structure of Alpine stonefly community assemblages across seven habitats using DNA-species

**DOI:** 10.1101/765578

**Authors:** Maribet Gamboa, Joeselle Serrana, Yasuhiro Takemon, Michael T. Monaghan, Kozo Watanabe

## Abstract

1. Stream ecosystems are spatially heterogeneous, with many different habitat patches distributed within a small area. The influence of this heterogeneity on the biodiversity of benthic insect communities is well documented; however, studies of the role of habitat heterogeneity in species coexistence and community assembly remain limited. Here, we investigated how habitat heterogeneity influences spatial structure (beta biodiversity) and phylogenetic structure (evolutionary processes) of benthic stonefly (Plecoptera, Insecta) communities.
2. We sampled 20 sites along two Alpine rivers, including seven habitats in four different reaches (headwaters, meandering, bar-braided floodplain, and lowland spring-fed). We identified 21 morphological species and delineated 52 DNA-species based on sequences from mitochondrial *cox1* and nuclear ITS markers. Using DNA-species, we first analysed the patterns of variation in richness, diversity, and community composition by quantifing the contribution of each of the four reaches and seven habitats to the overall DNA-species diversity using an additive partition of species diversity analysis and distance-based redundancy analysis. Using gene-tree phylogenies, we assessed whether environmental filtering led to the co-occurrence of DNA-species using a two-step analysis to find a phylogenetic signal.
3. The four reaches significantly contributed to DNA-species richness; with the meandering section having the highest contribution. However, we found that habitats had an effect on DNA-species diversity, where glide, riffle and, pool influenced the spatial structure of stonefly communities possibly due to a high species turnover.
4. Among the habitats, the pool showed significant phylogenetic clustering, suggesting high levels of evolutionary adaptation and strong habitat filtering. This community structure may be caused by long-term stability of the habitat and the similar requirements for co-occurring species.
5. Our study shows the importance of different habitats on the spatial and phylogenetic structure of stonefly community assemblies and sheds light on the habitat-specific diversity that may help improve conservation practices.

## 1. Introduction

Stream ecosystems provide some of the most heterogeneous features in landscapes, given the dynamic interaction between spatial elements (e.g., topography) and ecological process (e.g., hydrology) (Benda et al., 2004; Tockner & Stanford, 2002). This interaction creates a variety of different habitats throughout the longitudinal (upstream-downstream) gradient of the river, classified as lotic (running water) and lentic (standing water) (Dobson & Frid, 1998; Calow & Petts, 1996; Hauer & Lamberti, 1996). Within a river reach, habitat heterogeneity in a river channel defines environmental patches that impact aquatic community composition and species distribution (Brasil, Da Silva, Batista, Olivera, & Ramos, 2017; Dias-Silva, Cabetter, Juen, & De Marco, 2013); and creates environmental filtering to sort species with similar physiological requirements (Saito, Cianciaruso, Siqueira, Fonseca-Gessner, & Povoine, 2016; Mykra, Heino, &, Muotka, 2007; Webb, Ackerly, McPeek, & Donoghue, 2002). Aquatic insects are abundant, diverse, and broadly distributed (Lancaster & Downes, 2013). Many studies have reported a strong positive relationship between aquatic insects species diversity and spatial habitat heterogeneity (e.g. Astorga et al., 2014; Karaus, Larsen, Guillong, & Tockner, 2013; Arscott, Tockner, & Ward, 2005; Benda et al., 2004; Batista, Buss, Dorville, & Nessimian, 2001), however, these were only based on a a more localized habitat scale, and the relationship between assembly process on the river reach and habitat scales remains limited to a few studies (e.g. Astorga et al., 2014; Pastuchova, Lehotsky, & Greeskova, 2008).

Stream ecologists have reported a deeper insight on how communities are assembled, with the inclusion of genetic information (e.g. Hughes, Schmidt, & Finn, 2009). This allows understanding how evolutionary forces influence natural communities and how ecological processes influence genetic variation (Genung et al. 2011). Previous studies showed that local environmental heterogeneity may be the strongest driver of community assembly patterns using putative DNA-species diversity of aquatic insects along the river (Baselga et al., 2013; Finn, Bonada, Múrria, & Hughes, 2011). Recognizing DNA-species resulting by DNA sequences clustering analysis approach improve taxonomic workflows (Gamboa & Arrivillaga-Henriquez, 2019; Serrana, Miyake, Gamboa, & Watanabe, 2019; Gattoliatt, Rutschmann, Monaghan, & Sartori, 2018) by exposing cryptic species (closely related organisms with similar morphology but genetically distinct characters; Jackson, Battle, White, Pilgrim, Stein, Miller, & Sweeney, 2013) and enriching the biodiversity assessment of a region.

Among the methods to identify DNA-species, the Generalized Mixed Yule Coalescent model (GMYC; Pons et al., 2006) is a method commonly used. GMYC quantify the transition between inter- and intra-species branching events on an ultrametric tree (i.e., where the distance from the root of every branch is equal), using molecular markers, such as mitochondrial DNA (Fujisawa & Barraclough, 2013; Mynott, Webb, & Suter, 2011), or based on the congruency between mitochondrial and nuclear DNA (Rutschmann et al., 2016). However, despite widely used, the majority of the observations are based on DNA-species diversity changes along the river’s longitudinal gradient (Finn, Zamora-Muñoz, Múrria, Sáinz-Bariáin, & Alba-Tercedor, 2013; Gill, Harrington, Kondratieff, Zamudio, Poff, & Funk, 2013; Jackson et al., 2013), where the influence of other river dimensions (e.g., habitats), has only been described in few species, such as beetles (Ribera & Vogler, 2008) or caddisflies (Marten, Brandle, & Brandl, 2006). Therefore, investigating variations in the spatial patterns of the aquatic insect community composition using DNA-species in the river channel and their relationship with habitat heterogeneity will help gain a deeper insight on how the community assembly in aquatic ecosystems.

Studies on community assembly have not only focused on spatial patterns and composition changes analysis, a phylogenetic signal has been proposed as an additional approach to study community assembly (Violle et al., 2011; Webb et al., 2002). The basis of this approach is to compare the ecological traits (e.g. habitats) and phylogenetic structures in communities with those expected under null models (i.e., phylogenetic signal). Environmental filtering is thought to cause related species to co-occur (i.e., phylogenetic clustering) more than what is expected by chance, as closely-related species are assumed to share similar physiological requirements. By contrast, competitive exclusion (Cavender-Bares & Wilczek, 2003) and disturbance (Dinnage, 2009) causes the opposite pattern (phylogenetic overdispersion) as species compete for the same limiting resources. Although phylogenetic clustering and overdispersion have been observed in aquatic insect community assemblages at different geographical distances (Saito, Soininen, Fonseca-Gessner, & Siquiera, 2015a; Saito, Siquiera, & Fonseca-Gessner, 2015b; Saito et al., 2016), and in specific trait-based approaches (e.g., respiration strategy; Buchwalter, Jenkins, & Curtis, 2002), the assessment of the phylogenetic structure in different habitats have not been explored.

This study aims to investigate the role of seven habitats on the beta diversity and phylogenetic signal of the stonefly community composition using DNA-species. Stoneflies (Plecoptera) are aquatic insects that show more complex ecological (Lancaster & Downes, 2013) and evolutionary (Gamboa & Watanabe, 2019) responses to habitat change than other insects (Bojková, Rádková, Soldán, & Zahrádková, 2014), as they are sensitive to environmental conditions, such as low oxygen concentrations and high water temperatures (Prenda & Gallardo-Mayenco, 1999); leading to adaptive divergence among populations (Gamboa & Watanabe, 2019). We hypothesised that species distribution and community assembly are related to specific habitats and that habitat filtering acts on closely-related species with similar physiological requirements. By contrast, high levels of species diversity are expected to reduce community similarities among habitats and lead to phylogenetic overdispersion. Using two molecular markers (*cox1* and ITS) to delineate putative species, we aimed to: (1) quantify the contribution of four reaches and seven habitats to the overall DNA-species diversity, (2) determine the influence of habitats on community assembly, and (3) compare the phylogenetic structure of stoneflies communities in each of the seven habitats.

## 2. Methods

### 2.1 Study sites and sampling collection

We selected seven habitat types (Fig. 1); waterfall (vertical descent of watercourse), riffle (high turbulence water flow), glide (low turbulence water flow), pool (flowing water >1 m deep), wando (small-sized water in the head or tail on a sandy bar; as described by Ishida, Abekura, & Takemon, 2005), side-channel (small-sized water pass flowing on a sandy bar), and isolated pond (isolated still water habitat on the sandy bar). These were selected from over 20 sampling sites in two Alpine rivers in northeastern Italy. Fifteen of the sampling sites were located on the Tagliamento River (T01 to T15) and five were located on its major tributary, the Fella River (F01 to F05) (Fig. 2). The sampling sites were distributed across four types of reaches (4-1556 m a.s.l., Table 1); constrained headwaters, meandering (zigzagging river movement), bar-braided floodplain (convergent river bankfull width), and lowland spring-fed streams in the floodplain (Doering, Uehlinger, & Tockner, 2013). All reaches were separated at spatial scale without conflating each other along both rivers. All seven habitat types were sampled in <5 m at each site. Habitat diversity per site was measured using Simpsons diversity index with at least one observation per habitat and sampling site.

**Figure 1.**
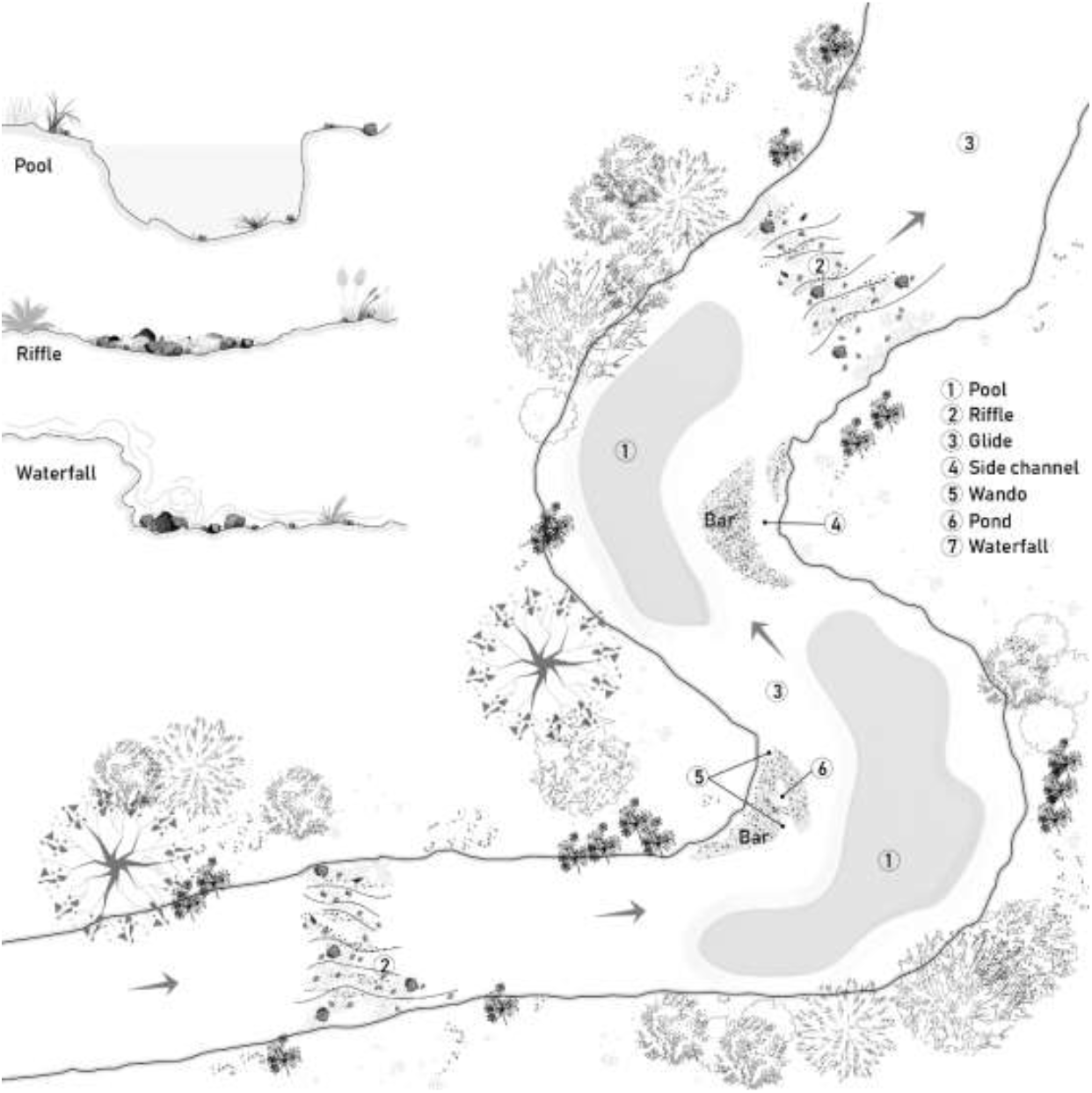
Habitats types sampled at each sampling site (Fig. 2) along the Tagliamento and Fella rivers. The arrows indicate the flow direction

**Figure 2.**
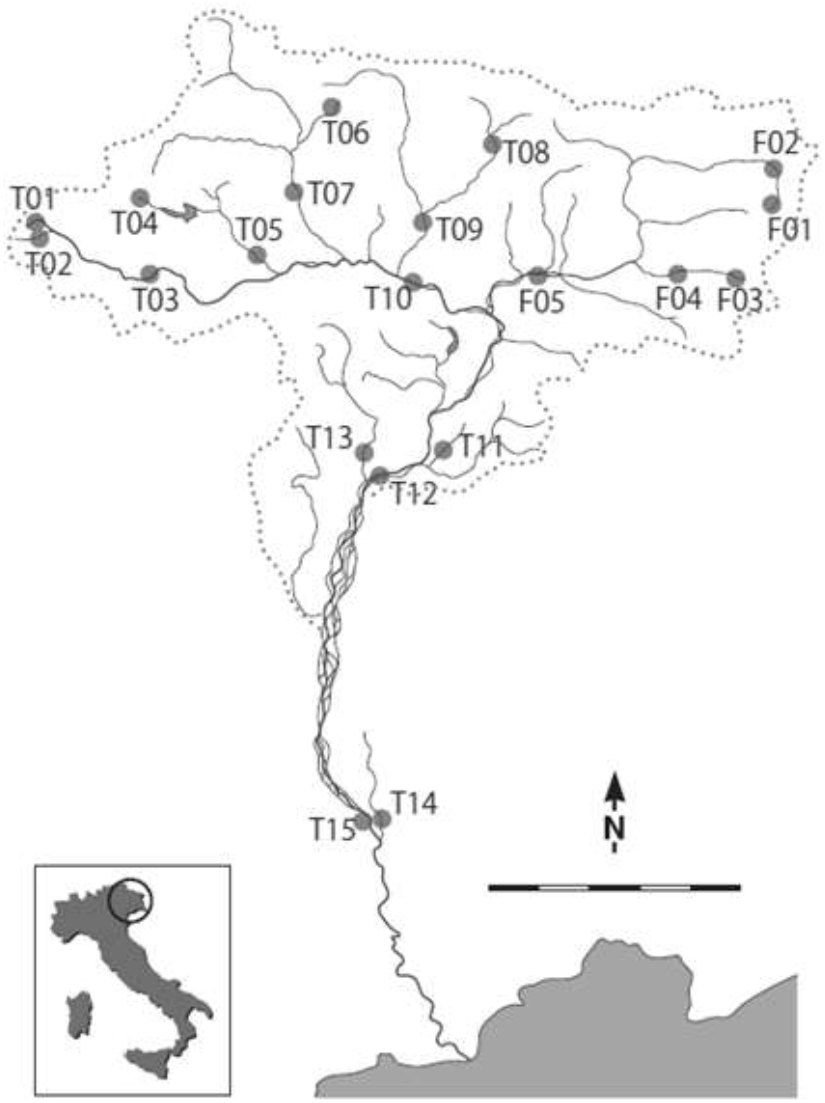
Sampling sites along the Tagliamento and Fella rivers. The rivers are located in the northeast region of Italy and sampling sites were coded using T1-15 (Tagliamento) and F1-05 (Fella)

**Table 1.** Sampling site characteristics, along with morphological and DNA-species richness in the Tagliamento (sites T) and Fella (sites F) rivers.

| Reaches | Sampling site | Altitude (m) | Habitat Simpsons diversity | Morpho-species | DNA species | Haplotypes |
| --- | --- | --- | --- | --- | --- | --- |
| Headwater | T01 | 1174 | 0.77 | 2 | 3 | 3 |
|  | T02 | 1064 | 0.77 | 3 | 4 | 6 |
|  | T04 | 1556 | 0.76 | 3 | 5 | 5 |
|  | T06 | 1346 | 0.78 | 8 | 10 | 10 |
|  | F01 | 1042 | 0.80 | 4 | 9 | 9 |
|  | F03 | 978 | 0.80 | 3 | 4 | 15 |
| Meandering | T05 | 486 | 0.76 | 6 | 10 | 12 |
|  | T08 | 625 | 0.81 | 5 | 15 | 12 |
|  | T13 | 183 | 0.80 | 5 | 10 | 13 |
|  | T15 | 5 | 0.81 | 2 | 5 | 2 |
|  | F02 | 769 | 0.80 | 4 | 10 | 13 |
|  | F04 | 488 | 0.76 | 2 | 2 | 2 |
| Bar-braided (floodplain) | T03 | 661 | 0.81 | 6 | 10 | 13 |
|  | T07 | 524 | 0.78 | 3 | 6 | 10 |
|  | T09 | 393 | 0.76 | 3 | 11 | 11 |
|  | T10 | 287 | 0.78 | 5 | 8 | 13 |
|  | T12 | 131 | 0.84 | 5 | 9 | 16 |
|  | F05 | 298 | 0.82 | 1 | 1 | 1 |
| Spring-fed | T11 | 170 | 0.55 | 4 | 11 | 15 |
|  | T14 | 4 | 0.55 | 1 | 1 | 3 |

We collected qualitative samples of stonefly nymphs using D-frame nets (250 μm mesh), spending one hour per site (total sampling time: 20 h). Between one and five replicates per habitat per site were collected (n = 188), where each replicate represented a different random location within each habitat sampled. Field sampling (all sites, and habitats) was carried out twice in summer 2009 (10 July-10 September) and once in early spring 2010 (24 March-15 April). All samples collected in the field were immediately immersed in >99 % ethanol and rinsed twice with fresh >99 % ethanol after being transported to the laboratory. Morphological species identification was conducted using a stereoscopic microscope (80 X) following the classifications of Consiglio (1980) and Fochetti and Tierno de Figueroa (2008).

### 2.2 DNA extraction and sequencing analysis

A total of 375 stonefly nymphs were collected (see Results). Genomic DNA was extracted from each nymph using DNeasy Blood and Tissue kits, according to the manufacturer’s instructions (Qiagen GmbH, Hilden, Germany). In all individuals, a 658 bp fragment of mitochondrial *cox1* was amplified using LCO-1490 and HCO-2198 primers (Folmer, Black, Hoeh, Lutz, & Vrijenhoek, 1994) with an annealing temperature of 48°C and 30 PCR cycles. The internal transcribed spacer (ITS) nuclear marker (average 930 bp) was amplified in a subset of 116 individuals chosen to represent all morphological taxonomic groups using ITS1 and ITS2 primers (McLain, Wesson, Oliver, & Collins, 1995) with an annealing temperature of 58°C and 40 PCR cycles. PCR products were purified using the QIAquick PCR Purification Kit (Qiagen) and sequenced in both directions using the aforementioned primers. *Cox1* sequences were analysed on a 3500xL automated sequencer (Applied Biosystems) and ITS sequences were analysed commercially by Eurofins - Operon, Tokyo. All sequencing data reported in this study have been deposited into GenBank (COI, MT482808 - MT483088; ITS, MT504421 - MT504536).

Forward and reverse reads were assembled and edited using CodonCode Aligner v 3.5 (Codon Code Corporation, Dedham, USA). Multiple sequence alignment was then performed using ClustalW (align.genome.jp; Larkin et al., 2007). Highly variable regions of ITS were removed using the Indel Module of SeqFIRE v 1.0.1 (https://omicstools.com/seqfire-tool; Ajawatanawong, Atkinson, Watson-Haigh, MacKenzie, & Baldauf, 2012) with the default settings. This reduced the ITS alignment length from 915-1016 bp to 790 bp. For both markers, identical sequences were collapsed using CleanCollapse v. 1.0.5 (https://sray.med.som.jhmi.edu/SCRoftware/CleanCollapse/). Sequences of mtDNA and nDNA markers were compared to the NCBI nucleotide database using blastn queries (http://blast.ncbi.nlm.nih.gov) to verify species identification (>90 and 80 % similarity for mtDNA and nDNA, respectively).

Genetic diversity for each marker was calculated as the number of polymorphic sites, nucleotide diversity (Kimura 2-parameter model), and the number of haplotypes using DnaSp v. 5.10 (Librado & Rozas, 2009).

### 2.3 Data analysis

#### 2.3.1 DNA-based species delimitation

Putative DNA-species were delineated using the General Mixed Yule Coalescent approach (GMYC; Fujisawa & Barraclough, 2013). First, ultrametric gene trees of *cox1* and ITS were constructed independently using BEAST v. 1.8.3 (Drummond, Suchard, Xie, & Rambaut, 2012) with a relaxed lognormal clock and a coalescent prior run for 50 million generations. The results were summarised with the Tree Annotator (BEAST package). A GMYC analysis was run using the splits package (Ezard, Fujisawa, & Barraclough, 2014) in R v. 3.3 (R Core Team, 2014). We used the single-threshold GMYC model based on Fujisawa and Barraclough (2013). The maximum likelihood of each GMYC model was tested using the likelihood ratio test against a one-species null model (where the entire tree is considered as a single coalescent). Outputs for *cox1* and ITS were compared and the results were combined to generate the list of DNA-species.

#### 2.3.2 Spatial structure of stonefly community assemblies

DNA-species diversity per reach and per habitat were calculated using the Shannon-Weaver and Simpsons diversity indices using the vegan package (Oksanen et al., 2012) in R. The contribution of each of the four reaches and seven habitat types to overall DNA-species diversity was quantified using an additive partition of species diversity analysis (Lande, 1996).

This analysis allows alpha and beta values to be simultaneously calculated using an abundance matrix organised hierarchically among habitats (alpha diversity, α) and reaches (beta diversity, β) (Gering, Crist, & Veech, 2003). Simpsons and Shannon-Weaver diversity were partitioned into alpha and beta diversity to observe whether the overall value was greater than the mean values (Lande, 1996). The statistical significance of each component was tested with a randomisation procedure (10,000 randomisations) using the adipart function of the vegan package in R. We then partitioned beta diversity into turnover (i.e., the replacement of species by other species in different habitats) and nestedness (i.e., species loss or gain between habitats) components (Baselga et al., 2013), and performed comparisons first, among habitats and then, habitats between each reaches using a Sorenson dissimilarity matrix and the betadisper function of the vegan package in R. We used a null model with 10,000 randomisations to test if the results of each component were greater than expected by chance.

#### 2.3.3 Habitat influence on community assembly

We performed a distance-based redundancy analysis (db-RDA; Legendre & Anderson, 1999) to quantify the influence of habitats on spatial community assemblages between sampling sites. We first conducted Principal Coordinates Analysis (PCoA) using a community dissimilarity matrix (the pairwise distance of DNA-species within each sampling site) calculated as the Bray-Curtis index. Then, we used the resulting eigenvalues as the “response” variable in the db-RDA models, while geographical distance, elevation (altitude, Table 1), reaches and habitat influence were used as explanatory variables. The geographical distance between each pair of sampling site was calculated as the Euclidean distance extracted from the geographical coordinates as proposed by Gamboa and Watanabe (2019) using the Vicenty Ellipsoid package (Kamey, 2013) in R. To analyse the contribution of the reaches and different habitats to the process of community assembly, we calculated a contribution diversity approach matrix as proposed by Lu, Wagner & Chen (2007). This approach calculated a differentiation coefficient to evaluate the distribution of DNA-species diversity in a given reach and habitat within the sampling sites. We obtained the best model by first including all explanatory variables (reaches, seven habitats, elevation, and geographical distance) then running nested models using the vegdist and capscale functions in the vegan package in R. The model was obtained using a permutation test (1000 permutations) and ANOVA to test each explanatory variable of the db-RDA model for statistical significance using the varpart function in vegan packge in R. After identifying the statistical significant model, we calculated the percentage of explained variance obtained by the db-RDA as suggested by Peres et al. (2006) using the ordistep function in vegan.

We analysed multivariate dispersion as a measure of beta diversity based on a community dissimilarity matrix using DNA-species to further investigate habitat influence on beta diversity. We calculated beta diversity as the distance to group centroid (the homogeneity of a community in a given habitat) based on a Bray-Curtis dissimilarity matrix. The community matrix consisted of DNA-species richness in each sampling site at both rivers. We also analysed the homogeneity of multivariate dispersions using a linear model in the betadisper function in the vegan R package with a permutation test (1000 permutations).

#### 2.3.4 Phylogenetic structure of the stonefly community

The phylogenetic relationship among species was estimated using MaximumLikelihood (ML) gene trees for *cox1* and ITS separately in PhyML 3.1 (Guindon & Gascuel, 2003), with the default settings under a GTR+I+G model as obtained in jModeltest 3.0 (Posada, 2008). Node support was determined by bootstrapping (10,000 times). We used sequences of Orthoptera sp. as outgroups in the *cox1* (HM381647) and ITS (KT440350) trees based on the reported phylogenetic relationship with stoneflies (Misof et al., 2014). Additional adult specimens (Gamboa & Monaghan, 2015) were added on the *cox1* tree to corroborate nymph species identification.

A congruency index (de Vienne, Giraud, & Martin, 2008) calculated using the web version of MAST (http://max2.ese.u-psud.fr/icong/index.help.html), indicated that *cox1* and ITS tree topologies were congruent (Icong = 1.9, p <0.05). Therefore, we concatenated the two markers using the FASconCAT-G perl script (https://github.com/PatrickKueck/FASconCAT-G). Using this matrix, we quantified the multisite phylogenetic similarities (i.e., similarity measures comparing more than two sampling sites) as proposed by Leprieur et al. (2012). We quantified multisite phylogenetic turnover, nestedness and phylo-beta diversity index (known as Sorensen similarity index) for every community by calculating the sum of the branch length in a phylogeny connecting all species in a community. We then observed the proportion of shared branch lengths between two sampling sites. We performed the analysis using the phylo.beta.multi function in the R package betapart (Baselga & Orne, 2012). We also performed a two-step analysis to check for a phylogenetic signal in stonefly communities within each habitat. First, we estimated niche conservatism (i.e., closely related species are ecologically similar and thus share similar habitats; Wiens et al., 2010) using Blomberg’s K-statistic (Blomberg, Garland, & Ives, 2003) and the phylogenetic independent contrast (PIC) test (Felsenstein, 1985) within each habitat. The Blomberg’s K test and PIC were implemented using the phytools package (Revell, 2012) and picante package (Kembel et al., 2010) in R. Both tests were run for 10,000 iterations to obtain the null distribution. If the p-values of the observed versus random variance were <0.05, we interpreted them as evidence of niche conservatism. Second, we calculated the Net Relatedness Index (NRI) and the Nearest Taxon Index (NTI) to measure the degree of phylogenetic clustering at the community level (Webb et al., 2002) within each habitat. NRI measures the phylogenetic dispersion of a community assemblage (community associated with the same habitat type) by comparing the observed mean pairwise phylogenetic distance between species in a community assemblage to the null model, while NTI measures the phylogenetic dispersion of a community assemblage by comparing the observed mean nearest phylogenetic neighbour distance between species in a community assemblage to the null model. Positive values of NRI and NTI indicate that community assemblages with similar habitat preferences are phylogenetically clustered (more closely related than expected), while negative values indicate phylogenetic overdispersion (more distantly related than expected). The NRI and NTI were calculated using the ses.mpd and ses.mntd functions, respectively, in R package picante, with 1000 replications.

## 3. Results

We identified a total of 21 morphological species in 12 stonefly genera (Table S1) based on collections of 5-40 individuals (mean = 19) and 1-8 morphological species (mean = 4) from each of the 20 sampling sites (Table 1). There were 184 *cox1* haplotypes from the 375 sequenced individuals and 87 ITS genotypes from the subset of 116 sequenced individuals. Samples were dominated by *Leuctra major* (50 and 40 % of the total haplotypes (*cox1*) and genotypes (ITS), respectively; Table S1). For both markers, the log-likelihood of the GMYC model at the threshold (*cox1* = 2514.495; ITS = 2819.385) was significantly higher than the null model for a single coalescent (*cox1* = 2400.932; ITS = 2701.875), as shown by the likelihood ratio test (p < 0.001). Both genes delimited 52 individual putative species, 38-59 species (95% CI) for *cox1* and 40-64 species (95% CI) for ITS (Table S1). The *cox1* gene was composed of 42 clusters and ten singletons, while ITS was composed of 31 clusters and 21 singletons. Intraspecific nucleotide diversity for *cox1* ranged from 0-13 % among putative DNA-species, being the highest value for *L*. *major* (Table S1). Fourteen of the 21 putative morpho-species were split into multiple DNA-species. All DNA-species occurred in more than one habitat, sampling site and, reach.

### 3.1 Spatial structure of stonefly community assemblies

Among the four reaches, the meandering contained the highest number of DNA-species (n = 52) (Table 1) and the highest Shannon-Weaver (0.89) and Simpson (0.6) diversities (Table S2). The single most species-rich site was T08 in the meandering section (n = 15 DNA-species) (Table 1). The headwater section contained the highest number of cryptic species (25 of 31 total cryptic DNA-species). Among the seven habitats, riffles contained the highest number of DNA-species (n = 31) followed by glides (n = 25) (Table 2 and S2). Riffles was the most DNA-species-rich habitat in the meandering section.

**Table 2.** Phylogenetic signal of stonefly communities in the seven habitats sampled in this study. Bold values indicate statistical significance (p <0.05). Positive values indicate phylogenetic clustering and negative values indicate overdispersion. N = total number of DNA-species per habitat, K = Blomberg’s K, PIC = phylogenetic independent contrasts, NRI = net relatedness index, NTI = nearest taxon index

|  | **N** | **K** | **PIC** | **NRI** | **NTI** |
| --- | --- | --- | --- | --- | --- |
| Waterfall | 21 | 2.30 | 2.41 | -1.29 | -0.79 |
| Riffle | 31 | 2.30 | 41.73 | 1.52 | **-1.85** |
| Glide | 25 | 2.71 | 26.47 | 0.11 | -0.87 |
| Pool | 10 | **4.03** | **76.80** | **2.53** | **1.79** |
| Wando | 6 | 1.73 | 10.89 | **-6.44** | -1.26 |
| Side channel | 4 | 1.25 | 64.83 | **-2.06** | -1.06 |
| Isolated Pond | 17 | 2.09 | 16.59 | 0.15 | -0.82 |

The relative contribution of the four reaches and seven habitats to overall DNA-species diversity showed that reaches were the most important contributor for richness (57 %, p < 0.001, Fig. 3), while habitats were the most important contibutor for species diversity (Shannon-Weaver index; 88 %, p < 0.001). DNA-species turnover among habitats was significantly higher (0.60) than species nestedness (0.16) to overall beta diversity (0.79) (p = 0.0016), similarly high species turnover (0.58) was found among habitats between reaches (p = 0.0020).

**Figure 3.**
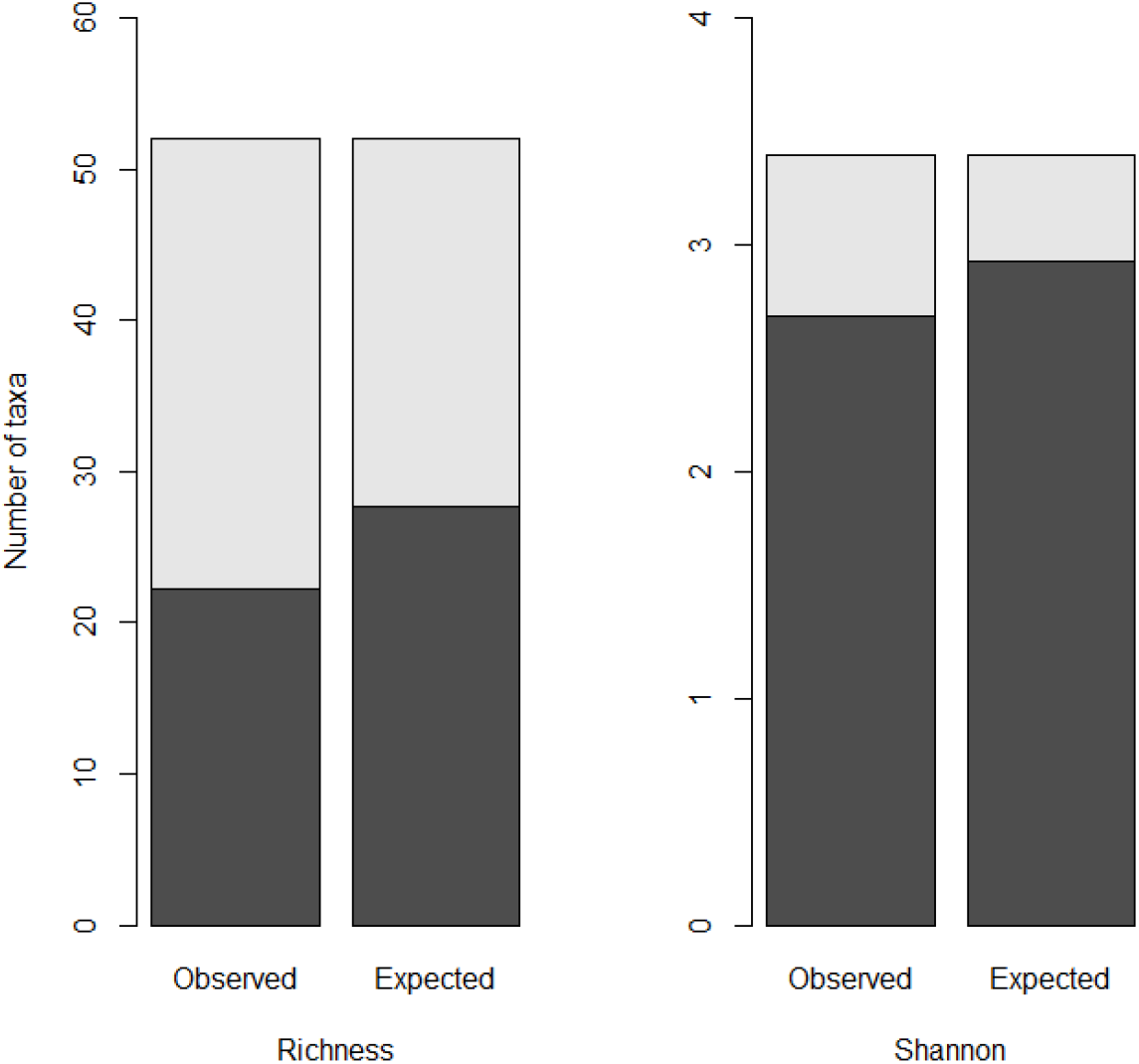
Additive diversity partition of species richness and Shannon-Weaver index (based on DNA-species) into observed and expected hierarchical components of reaches (light grey) and habitats (dark grey). Both partition analyses were significant (<0.001) based on 10,000 randomisations. Note that the plots have different scales

Spatial variation in DNA-species was only found to be significant in three habitats: riffle, glide, and pool (70% of total variation, p <0.05, Fig. 4). Among the seven habitats, the riffle and glide had the highest diversity dissimilarity (F = 1.43, p <0.05, Fig. 5), suggesting that the community in these two habitats were less similar among each other that in the remaining habitats.

**Figure 4.**
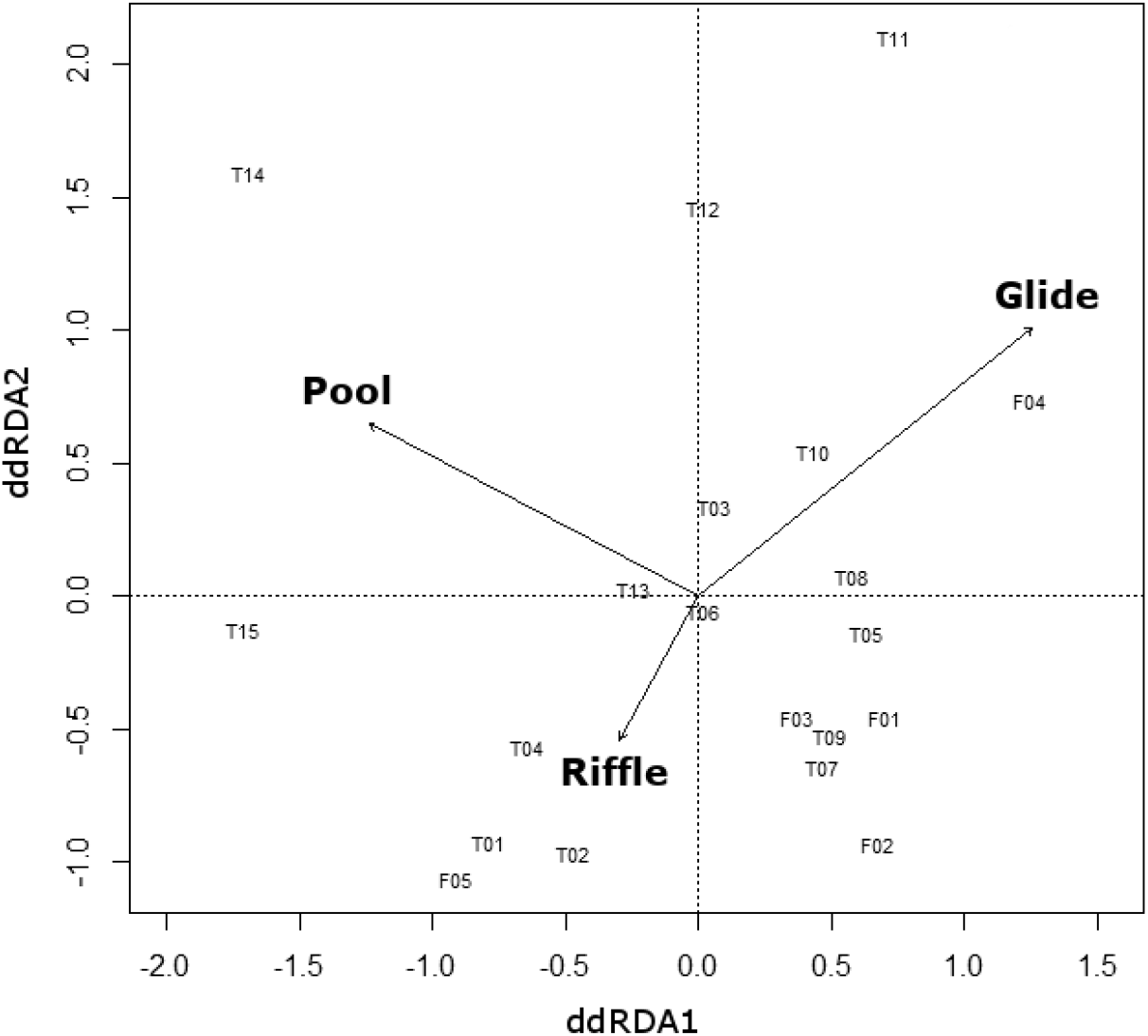
Distance based-redundancy analysis (dbRDA) of stonefly DNA-species among 20 sampling sites displaying the influence of habitats on community structure. T = Tagliamento river, F = Fella river. Three habitats (pool, glide, and riffle) explained the highest proportion of the total variation

**Figure 5.**
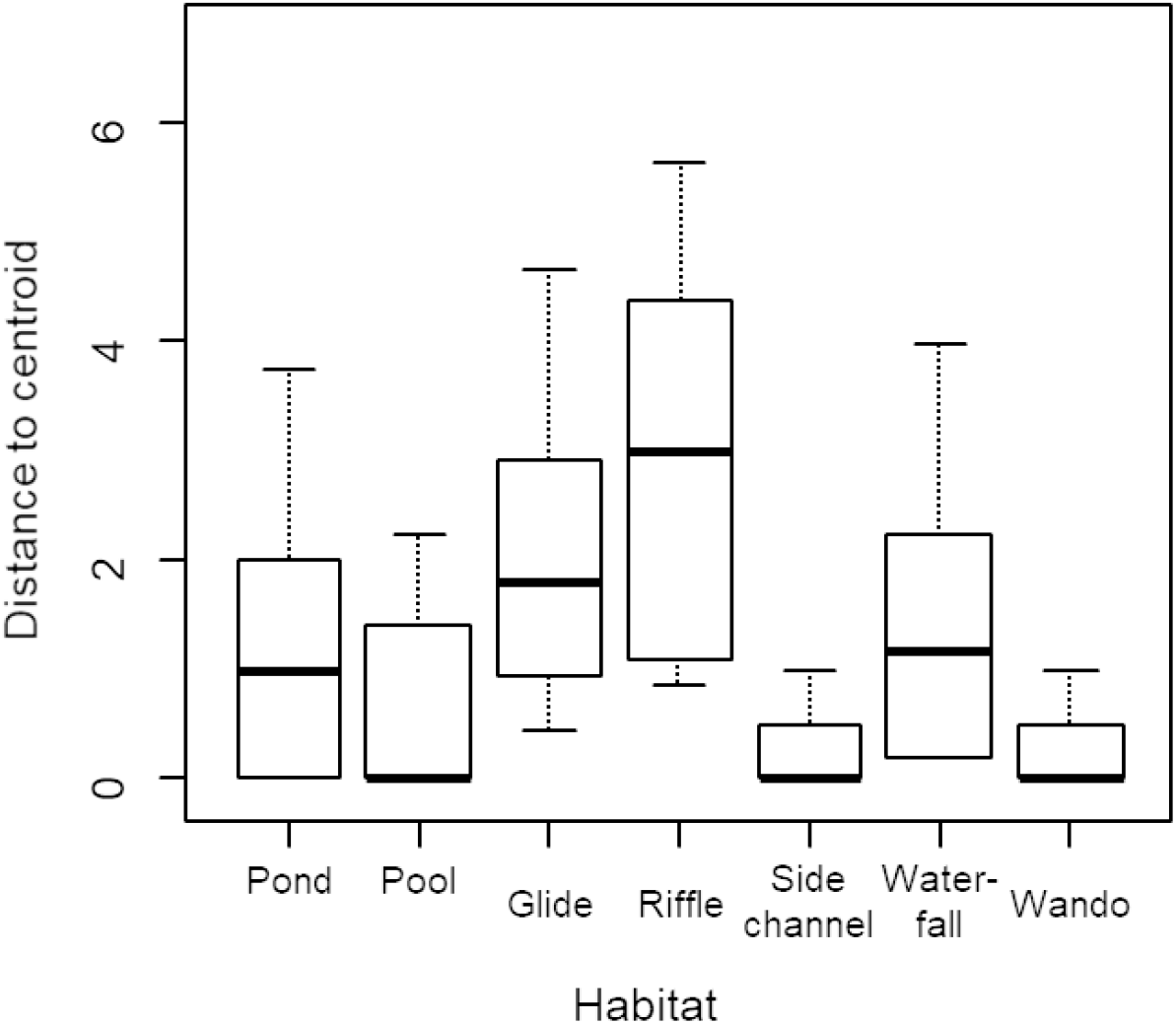
Beta diversity of stonefly communities in each of the seven habitats, based on the distance of the values for each habitat to their centroid (community homogeneity in a given habitat, see methods section 2.3.3). The bold line represents the median, while the bottom and top of the box represent lower and upper quartiles, respectively

### 3.2 Phylogenetic structure of stonefly community

The mtDNA phylogeny recovered all recognised morphospecies as clades; however, 11 morphospecies were composed of multiple lineages. The multiple lineages were found within the same reaches. These lineages likely reflect cryptic species, concurring with GMYC results. The nuclear ITS tree showed congruence with the *cox1* tree, except those of *Nemoura cinerea* and *N. mortini* which were located in the same clade as *Capnia nigra* and *C. vidua* (Fig. S1).

The stonefly community phylogenetic diversity among sampling sites showed that species turnover (0.927) contributed more than nestedness (0.055) to beta diversity (0.98) (p < 0.005). Pool habitats showed significant niche conservatism and phylogenetic clustering based on the two-step analysis of phylogenetic signal (Blomberg’s K test, PIC, NRI and, NTI, all *p* < 0.05; Table 2), suggesting high similarity among closely-related species within this habitat. Additionally, significant negative values for NRI and NTI indices were reported (*p* < 0.05, Table 2) in the wando and side channel for NRI, and riffle for NTI, indicating that the community assemblage presents phylogenetic overdispersion.

## 4. Discussion

Community assemblages within the stream are strongly related to habitat heterogeneity within a given area (Astorga et al., 2014). We observed that different habitat types played a role in Alpine stoneflies DNA-species diversity changes, their spatial structure, and their evolution.

In this study, the habitats had the most important role in DNA-species diversity (diversity partition analysis, Shannon index), concurring with previous studies of aquatic invertebrates (Astorga et al., 2014; Finn & LeRoy Poff, 2011; Finn, Theobald, Black, & Poff, 2006; Finn et al., 2011; Ribera & Vogler, 2008; Marten et al., 2006). Additionally, our results showed a high species turnover among habitats (species turnover = 0.60). A high species turnover is associated with poor dispersal ability (Thompson & Townsend, 2006) commonly associated with spatial restrictions due to niche heterogeneity (Heino et al., 2015; Astorga et al., 2014; Finn & LeRoy Poff, 2011). Therefore, our results suggest that non-random migration is likely to occur among habitats, probably caused by environmental filtering or spatial restrictions which may affect the dispersal ability of the species. Stoneflies have relatively poor dispersal abilities within stream reaches (Consiglio, 1980; Fochetti & Tierno de Figueroa, 2008), and tend to avoid migration among habitats due to predator pressure (Tiziano, Fenoglio, Lopez-Rodriguez, Tierno de Figueroa, Grenna, & Cucco, 2010). The habitat heterogeneity along the Tagliamento and Fella rivers could potentially reduce the intra-stream stonefly dispersal, and might explain the large number of cryptic species found (GMYC and phylogenetic analyses). However, high species turnover is also associated with sampling bias. Non-common species among habitats could be a consequence of an unequal sampling effort or habitat availability. In this study, habitat availability was similar across most sampling sites (average = 0.75, range = 0.55-0.84, Table 1) and despite a similar sampling effort at every site, some species were not detected. Therefore, future studies will need to increase sampling effort to estimate if sampling bias could impact our interpretations of species turnover.

Among the seven habitats, the riffle, glide, and pool had significant impacts on the stonefly community structure (according to the db-RDA analysis) and two habitats (riffle and glide) had significantly higher species diversity (according to the multivariate dispersion analysis). The riffle and glide habitats characterised by a high and low turbulence flow, respectively; both play an essential role in determining habitat suitability for many species of aquatic insects (Lancaster & Downes, 2013; Benda et al., 2004), particularly stoneflies (Lancaster & Downes, 2013; Batista et al., 2001). The dynamic nature of riffle and glide habitats influence density-dependent local competition and high organisms migration with the turbulent flow, leading to high levels of biodiversity (Hughes et al., 2009; Arscott et al., 2005; Batista et al., 2001), as observed in stoneflies (Prenda & Gallardo-Mayenco, 1999). Despite our results, pools (a deep-high river section with a slow/nonexistent velocity) have been observed to be weekly correlated with aquatic insect community diversity (Herrera-Vasquez, 2008) due to the specialised taxa living in this stable habitat (Pastuchova, Lehotsky, & Greeskova, 2008).

Surprisingly, we also found that the pool habitat led to a phylogenetic clustering of the stonefly community according to the four indices used in this study (Blomberg’s K, PIC, NRI, NTI). Long-term adaptation (Lososova et al., 2015), dispersal limitations (Saito et al., 2015a, 2015b), colonisation history (Aizen, Gleiser, Sabatino, Gilarranz, Bascompte, & Verdu, 2015), and low resource competition (Webb et al., 2002) are all associated with phylogenetic clustering, particularly in habitats linked with long-term stability. Pools are rarely affected by physical perturbations (Buffington, Lisle, Woodsmith, & Hilton, 2002), and they may persist despite high river flow (Calow & Petts, 1996), discharge (Rolls, Leigh, & Sheldon, 2012) and drought (Lake, 2003). Therefore, habitats with low levels of perturbation enhanced niche conservatism, clustering and, evolutionary adaptation of stonefly communities in Alpine rivers. However, a greater competitive asymmetry (i.e., an unequal resource division) among distant phylogenetic relatives and facilitation among close phylogenetic relatives can also cause phylogenetic clustering (Mayfield & Levine, 2010; Sargent & Ackerly, 2008). Future studies should address this concern by studying the distribution of resources (i.e., food) among habitats, to obtain a clearer understanding of the influences on phylogenetic clustering of co-occurring species.

In riffle, wando and side channel habitats, a phylogenetic overdispersion pattern was detected, which is commonly observed in aquatic insects (Saito et al., 2016; Violle et al., 2011). Efficient colonisation and high dispersal patterns are thought to be driving factors for the phylogenetic overdispersion of several species groups, such as birds (Sobral & Cianciaruso, 2016), mammals (Cardillo, Gittleman, & Purvis, 2008) and insects (Violle et al., 2011). These characteristics have been rarely documented in stoneflies (e.g., *Leutra ferruginea*, Macneale, Peckarsky, & Likens, 2005); however, stonefly nymph dispersal is driven by intra-stream drift (Stewart & Stark, 2008). Phylogenetic overdispersion is common in highly-disturbed habitats which tend to preserve high phylogenetic diversity (Xiu et al., 2017). Habitats with high-turbulent water flow (riffle) and high physical perturbation such as sandy bar habitats (wando and side-channel) in the Tagliamento river show a 62 % habitat turnover (Tockner et al., 2003) mainly driven by two annual flooding events (Ward et al., 1999). Therefore, the phylogenetic overdispersion of stonefly communities at riffle, wando, and side-channel habitats is likely driven by the high level of disturbance or possible natural flow disturbances in these habitats.

Overall, the spatial and the phylogenetic structure of stonefly community assemblages in the Tagliamento and Fella rivers were promoted by the availability and stability of habitats. Among the four reaches, meandering reach harboured the largest number of DNA-species (n = 52) and was highly diverse among habitats (Shannon-Weaver index: 0.89). Meandering stream sections are highly complex morphodynamic sections of the river supporting high benthic invertebrate diversity, given the suitability and resilience of their habitats within fluvial ecosystems (Garcia, Schnauder, & Push, 2012). However, the headwaters exhibited a high number of cryptic species (78% of the total DNA-species richness), as previously shown by Jackson et al. (2013), Múrria et al. (2013), Finn et al. (2011), and Hughes et al. (2009). This tendency is due to lower canalisation, loss of connectivity with the stream network, and low levels of anthropogenic disturbance (Finn & LeRoy Poff, 2011; Finn et al., 2011). Cryptic species are often observed in stoneflies (e.g., Viteck, Vincon, Graf, & Pauls, 2017), possibly due to introgression (Boumans, Hogner, Brittain, & Johnsen, 2017) or hybridisation (Young, Smith, Pilgrim, Fairchlid, & Schwartz, 2019) which remain unresolved species. Headwaters in the Tagliamento and Fella rivers have narrow valleys and gorges with low sinuosity and confinement (Tockner et al., 2003); therefore, this reach enhances intraspecific genetic differences among stoneflies species. Both meandering and headwater reaches strongly influenced the maintenance of stonefly biodiversity in alpine rivers; therefore, their environmental protection should be a priority in achieving conservation practices. We recommend that future studies should take into consideration the river characteristics as geomorphological information and perform additional comparisons among habitats within a sampling location to understand how the habitats play a role in our results.

In conclusion, we demonstrated that different habitats played an essential role in the spatial and phylogenetic structure of stonefly community assemblages in alpine regions. Community diversity was related with habitats with turbulent water flow, while phylogenetic clustering was related with habitats with long-term stability. Our results highlight the importance of molecular studies on quantitative measures of biodiversity in given area and the essential role of habitats in a region, which is particularly important to conserve community assemblages with threatened species or within threatened habitats/landscapes (Esteban & Finlay, 2010). Despite the differences among habitats between reaches, a phylogenetic signal could be detected in the pool habitats. This signal could be an indication that habitats strongly influence the biodiversity of a region. Our results could be further improved by adding information about habitat geomorphology and environmental factors. Therefore, we recommend that future studies should include and address habitat characteristics, as well as comparisons among habitats between reaches to further understand species diversity changes.

## Acknowldedgements

We thank Paul Schmidt-Yáñez for his assistance in the field, Micanaldo Francisco for his assistance with the figures, and anonymous reviewers for their constructive comments. MG was supported by the German Academic Exchange Service (DAAD) fellowship (A/09/94531) and the Japan Society of the Promotion of Science Postdoctoral Fellowship (PU17908). KW was supported by a European Union Marie-Curie International Incoming Fellowship (PIIF-GA- 2009-237026). MTM was partially supported by a Japan Society for the Promotion of Science (JSPS) Fellowship (L-15543). This research was supported by the JSPS (Grant Numbers: 24254003, 17H01666), the Sumitomo Electric Industries Group Corporate Social Responsibility Foundation, the German Academic Exchange Service (DAAD, Programm Projektbezogener Personenaustausch Japan, project 57402018) and the Research Unit Program of Ehime University.

## Data availability statement

The data that support the findings of this study are available from the corresponding author upon reasonable request.

## Supplementary Material

**TABLE S1.** DNA-species list for all sampling sites. Present sampling sites: number of sampling sites were DNA-species were found. Present habitats: different types of habitats were each DNA-species was found.

| Morphospecies | Total<br>n | DNA-species | Haplotypes | Present<br>sampling<br>sites | Present<br>habitats | Genetic<br>Diversity (%) | Altitude<br>range (m) |
| --- | --- | --- | --- | --- | --- | --- | --- |
| Systellognatha |  |  |  |  |  |  |  |
| Perlodidae |  |  |  |  |  |  |  |
| *Isoperla* cf. *andreinii* | 2 | 1 | 1 | 1 | 1 | 0 | 1346 |
| *Isoperla grammatica* | 15 | 2 | 9 | 5 | 4 | 2 | 486-1346 |
| *Besdolus* sp. | 28 | 3 | 14 | 7 | 5 | 3.5 | 183-1556 |
| *Perlodes microcephalus* | 16 | 2 | 10 | 6 | 2 | 2 | 287-769 |
| *Perlodes intricatus* | 4 | 1 | 3 | 3 | 2 | 0 | 287-625 |
| Perlidae |  |  |  |  |  |  |  |
| *Dinocras cephalotes* | 13 | 2 | 8 | 4 | 2 | 2.6 | 4-183 |
| *Perla marginata* | 24 | 2 | 15 | 6 | 5 | 6.5 | 131-661 |
| *Perla grandis* | 4 | 2 | 3 | 2 | 1 | 3 | 183-625 |
| Chloroperlidae |  |  |  |  |  |  |  |
| *Chloroperla susemicheli* | 13 | 5 | 9 | 2 | 5 | 9.5 | 287-661 |
| Euholognatha |  |  |  |  |  |  |  |
| Capniidae |  |  |  |  |  |  |  |
| *Capnia nigra* | 17 | 3 | 9 | 2 | 2 | 7 | 131-170 |
| *Capnia vidua* | 8 | 1 | 2 | 1 | 2 | 0 | 170 |
| Nemouroidae |  |  |  |  |  |  |  |
| *Nemoura mortoni* | 8 | 3 | 5 | 4 | 4 | 8.2 | 486-1346 |
| *Nemoura cinerea* | 10 | 1 | 2 | 1 | 2 | 0 | 1346 |
| *Amphinemoura* sp. | 2 | 1 | 1 | 1 | 1 | 0 | 524 |
| *Protonemoura* cf.<br>*nimborum* | 3 | 2 | 3 | 3 | 4 | 5.2 | 393-1174 |
| *Protonemoura*<br>*nimborella* | 7 | 1 | 1 | 1 | 1 | 0 | 1556 |
| *Protonemoura nitida* | 6 | 2 | 4 | 1 | 2 | 1.3 | 1346 |
| Leuctricidae |  |  |  |  |  |  |  |
| *Leuctra fusca* | 25 | 7 | 12 | 6 | 3 | 3.7 | 131-287 |
| *Leuctra braueri* | 10 | 2 | 6 | 2 | 3 | 4.2 | 978-1346 |
| *Leuctra major* | 91 | 8 | 64 | 18 | 7 | 13.1 | 5-1556 |
| Taenioterygidae |  |  |  |  |  |  |  |
| *Brachyptera risi* | 6 | 1 | 3 | 1 | 2 | 0 | 131 |

**TABLE S2.** DNA-species richness among habitats per sampling site

| Reaches | Sampling sites | Waterfall | Riffle | Glide | Pool | Wando | Side-channel | Isolated Pond | Shannon diversity | Simpson diversity |
| --- | --- | --- | --- | --- | --- | --- | --- | --- | --- | --- |
| Headwater | T01 | 0 | 0 | 3 | 0 | 0 | 0 | 0 | 0.048602 | 7.68E-05 |
|  | T02 | 4 | 2 | 1 | 0 | 0 | 0 | 0 | 0.092222 | 0.000538 |
|  | T04 | 3 | 3 | 1 | 0 | 0 | 0 | 3 | 0.119007 | 0.001152 |
|  | T06 | 4 | 10 | 4 | 0 | 0 | 1 | 2 | 0.19427 | 0.005376 |
|  | F01 | 13 | 7 | 1 | 0 | 2 | 0 | 3 | 0.220693 | 0.008321 |
|  | F03 | 0 | 5 | 7 | 0 | 0 | 0 | 0 | 0.134995 | 0.00169 |
| Meandering | T05 | 3 | 4 | 2 | 3 | 0 | 0 | 4 | 0.163554 | 0.003072 |
|  | T08 | 1 | 17 | 1 | 0 | 6 | 0 | 2 | 0.225542 | 0.008986 |
|  | T13 | 5 | 6 | 8 | 0 | 2 | 1 | 3 | 0.215707 | 0.00768 |
|  | T15 | 0 | 0 | 5 | 1 | 0 | 0 | 0 | 0.082351 | 0.000384 |
|  | F02 | 1 | 11 | 1 | 0 | 3 | 0 | 1 | 0.170096 | 0.003482 |
|  | F04 | 0 | 1 | 0 | 1 | 0 | 0 | 0 | 0.035297 | 2.56E-05 |
| Bar-braided (floodplain) | T03 | 6 | 8 | 1 | 2 | 0 | 1 | 3 | 0.19427 | 0.005376 |
|  | T07 | 0 | 2 | 4 | 1 | 1 | 0 | 1 | 0.110493 | 0.000922 |
|  | T09 | 2 | 7 | 7 | 0 | 0 | 0 | 0 | 0.163554 | 0.003072 |
|  | T10 | 2 | 4 | 4 | 2 | 0 | 0 | 6 | 0.176427 | 0.003917 |
|  | T12 | 1 | 6 | 8 | 3 | 0 | 1 | 1 | 0.188504 | 0.004864 |
|  | F05 | 0 | 0 | 0 | 0 | 0 | 2 | 0 | 0.035297 | 2.56E-05 |
| Spring-fed | T11 | 2 | 5 | 4 | 8 | 0 | 0 | 0 | 0.18256 | 0.004378 |
|  | T14 | 0 | 1 | 0 | 2 | 0 | 0 | 0 | 0.048602 | 7.68E-05 |
| Total |  | 47 | 99 | 62 | 23 | 14 | 6 | 29 | 2.802043 | 0.936585 |

## List of supplementary figures

**FIGURE S1.**
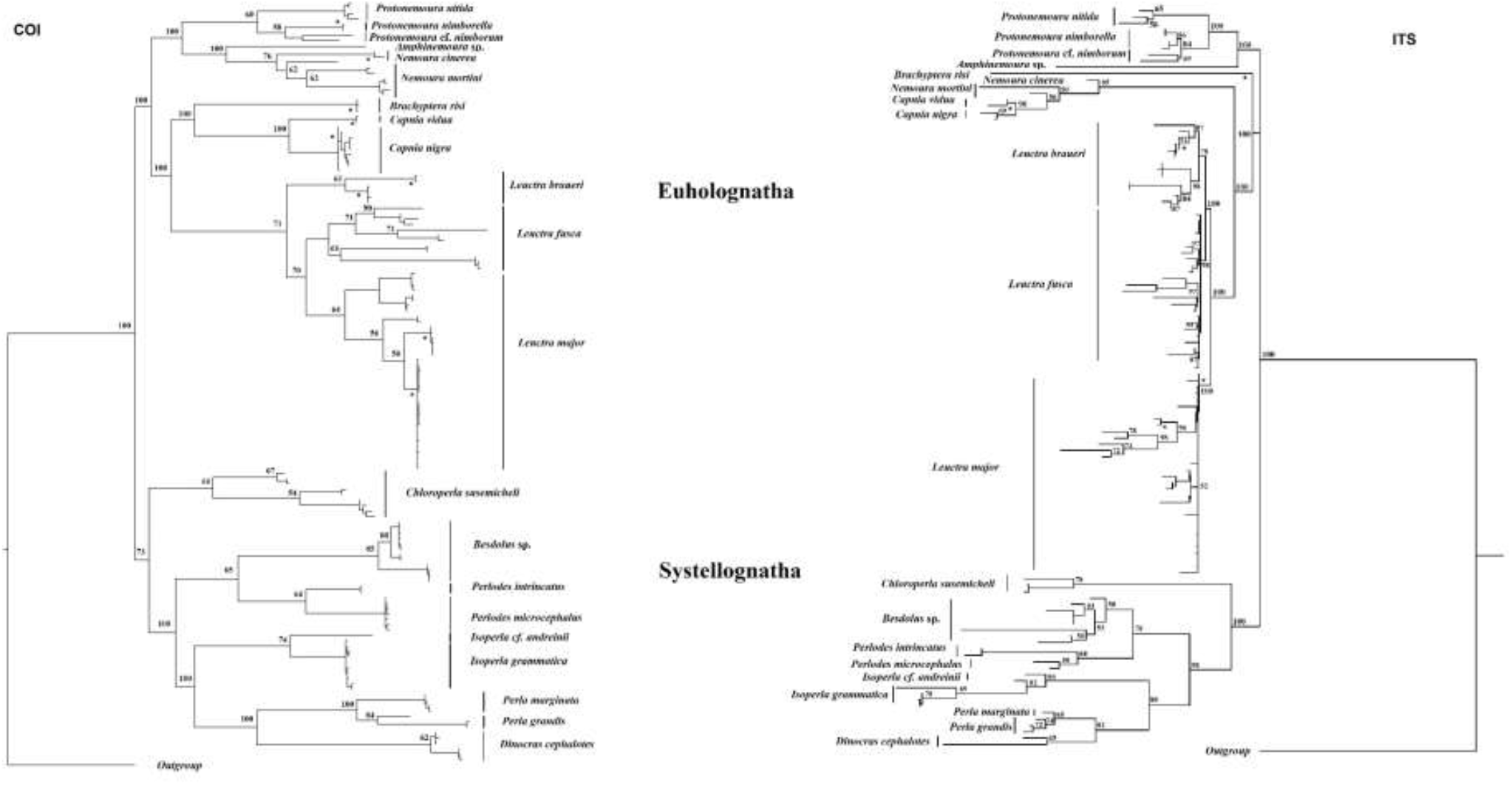
Maximum likelihood phylogenetic trees of stonefly species located along theTagliamento and Fella rivers. The *cox1* (mtDNA) tree is shown on the left and ITS (nDNA) on the right. The numbers at each node indicate the likelihood bootstrap support values. Asterisks show the adult specimen position. A subdivision in two morphological groups was observed, Euholognatha and Systellogna

## References

Aizen, M. A., Gleiser, G., Sabatino, M., Gilarranz, L. J., Bascompte, J., & Verdu, M. (2015). The phylogenetic structure of plant-pollinator networks increases with habitat size and isolation. Ecology Letters, 19, 29–36.

Ajawatanawong, P., Atkinson, G. C., Watson-Haigh, N. S., MacKenzie, B., & Baldauf, S. L. (2012). SeqFIRE: a web application for automated extraction of indel regions and conserved blocks from protein multiple sequence alignments. Nucleic Acids Research, 40, 340–347.

Arscott, D. B., Tockner, K., & Ward, J. V. (2005). Lateral organization of aquatic invertebrates along the corridor of a braided floodplain river. Journal of North American Benthological Society, 24, 934–954.

Astorga, A., Death, R., Death, F., Paavola, R., Chakraborty, M., & Muotka, T. (2014). Habitat heterogeneity drives the geographical distribution of beta diversity: the case of New Zealand stream invertebrates. Ecology and Evolution, 4, 2693–2702.

Baselga, A., & Orme, C. D. L. (2012). Betapart: and R package for the study of beta diversity. Methods in Ecology and Evolution, 3, 808–812.

Baselga, A., Fujisawa, T., Crampton-Platt, A., Bergsten, J., Foster, P. G., Monaghan, M. T., & Vogler, A. P. (2013). Whole-community DNA barcoding reveals a spatio-temporal continuum of biodiversity at species and genetics levels. Nature Communications, 4, 10.1038.

Batista, D. F., Buss, D. F., Dorville, L. F. M., & Nessimian, J. L. (2001). Diversity and habitat preference of aquatic insects along the longitudinal gradient of the Mace river basin, Rio de Janeiro, Brazil. Revista Brasileira de Biologia, 6, 249–258.

Benda, L., Poff, N. L., Miller, D., Dunne, T., Reeves, G., Pess, G., & Pollock, M. (2004). The network dynamics hypothesis: how channel networks structure riverine habitats. BioScience, 54, 413–427.

Blomberg, S. P., Garland, T., & Ives, A. R. (2003). Testing for phylogenetic signal in comparative data: behavioral traits are more labile. Evolution, 57, 717–745.

Bojková, J., Rádková, V., Soldán, T., & Zahrádková, S. (2014). Trends in species diversity of lotic stoneflies (Plecoptera) in the Czech Republic over five decades. Insect Conservation and Diversity, 7, 252–262.

Boumans, L., Hogner, S., Brittain, J., & Johnsen, A. (2017). Ecological speciation by temporal isolation in a population of the stonefly *Leuctra hippopus* (Plecoptera, Leuctridae). Ecology and Evolution, 7, 1635–1649.

Brasil, L. S., Da Silva, N. F., Batista, J. D., Olivera, B., & Ramos, H. S. (2017). Aquatic insects in organic and inorganic hábitats in the streams on the Central Brazilian savanna. Revista Colombiana de Entomologia, 43, 286–291.

Buchwalter, D. B., Jenkins, J. J., & Curtis, L. R. (2002) Respiratory strategy is a major determinant of [^3^H]water and [^14^C]chlorpyrifos uptake in aquatic insects. Canadian Journal of Fisheries and Aquatic Science, 59, 1315–1322.

Buffington, J. M., Lisle, T. E., Woodsmith, R. D., & Hilton, S. (2002). Controls on the size and occurrence of pools in coarse-grained forest rivers. River Research and Applications, 18, 507–531.

Calow, P. P., & Petts, G. E. (1996). The Rivers Handbook: Hydrological and Ecological Principles. Blackwell Science LTD, Victoria, Austria.

Cardillo, M., Gittleman, J. L., & Purvis, A. (2008). Global patterns in the phylogenetic structure of island mammal assemblages. Proceedings of the Royal Society B: Biological Science, 275, 1549–1556.

Cavender-Bares, J., & Wilczek, A. (2003). Integrating micro and macroevolutionary processes in community ecology. Ecology, 84, 592–597.

Consiglio, C. (1980). Plecoptteri. Consiglio Nazionale delle Ricerche AQ/1/77, Verona, Italy.

Dobson, M. & Frid, C. (1998). Ecology of Aquatic Systems. Longman, Harlow, UK.

Doering, M., Uehlinger, U., & Tockner, K. (2013). Vertical hydrological exchange, and ecosystem properties and processes at two spatial scales along a floodplain river (Tagliamento, Italy). Freshwater Science, 32, 12–25.

Dias-Silva, K., Cabetter, H. S. R., Juen, L., & De Marco JR, P. (2010). The influence of habitat integrity and physical-chemical water variables on the structure of aquatic and semi-aquatic Heteroptera. Zoologia, 27, 918–930.

Dinnage, R. (2009). Disturbance alters the phylogenetic composition and tructure of plant communities in an old field system. PLos ONE, 4, e7071.

Drummond, A. J., Suchard, M. A., Xie, D., & Rambaut, A. (2012). Bayesian phylogenetics with BEAUti and the BEAST 1.7. Molecular Biology and Evolution, 29, 1969–1973.

Esteban, G. F., & Finlay, B. J., (2010). Conservation work is incomplete without cryptic biodiversity. Nature, 463, 293.

Ezard, T., Fujisawa, T., & Barraclough, T. (2014). Splits: SPecies llmits. Retrieved from http://R-Forge.R-project.org/projects/splits/

Felsenstein, J. (1985). Phylogenies and the comparative method. American Naturalist, 125, 1–15.

Finn, D. S., & LeRoy Poff, N. (2011). Examining spatial concordance of genetic and species diversity patterns to evaluate the role of dispersal limitation in structuring headwater metacommunities. Journal of North American Benthological Society, 30, 273–283.

Finn, D. S., Bonada, N., Múrria, C., & Hughes, J. M. (2011). Small but mighty: headwaters are vital to stream network biodiversity at two levels of organization. Journal of the North American Benthological Society, 30, 963–980.

Finn, D. S., Theobald, D. M., Black, W. C., & Poff, N. L. (2006). Spatial population genetic structure and limited dispersal in a Rocky Mountain alpine stream insect. Molecular Ecology, 15, 3553–3566.

Finn, D. S., Zamora-Muñoz, C., Múrria, C., Sáinz-Bariáin, M., & Alba-Tercedor, J. (2013). Evidence from recently deglaciated mountain ranges that *Baetis alpinus* (Ephemeroptera) could lose significant genetic diversity as alpine glaciers disappear. Freshwater Science, 33, 207–2016.

Fochetti, R., & Tierno de Figueroa, J. M. (2008). Plecoptera. Fauna D’ Italia, Calderini, Italy.

Folmer, O., Black, M., Hoeh, W., Lutz, R., & Vrijenhoek, R. (1994). DNA primers for amplification of mitochondrial cytochrome c oxidase subunit I from diverse metazoan invertebrates. Molecular Marine Biology and Biotechnology, 3, 294–297.

Fujisawa, T., & Barraclough, T. G. (2013). Delimiting species using single-locus data and the generalized mixed yule coalescent approach: a revised method and evaluation on simulated data sets. Systematic Biology, 62, 707–724.

Garcia, X.-F., Schnauder, I., & Push, M. T. (2012). Complex hydromorphology of meanders can support benthic invertebrate diversity in rivers. Hydrobiologia, 685, 49–68.

Gamboa, M. & Arrivillaga-Henriquez, J. (2019). Biochemical and molecular differentiation of *Anacroneuria* species (Plecoptera, Insecta) in Andean National Park, Venezuela. Systematics and Biodiversity, 17, 669–678.

Gamboa, M. & Monaghan, M.T. (2015). Association of adult female and male stoneflies (Plecoptera) from an Alpine river using wing morphometrics and mitochondrial DNA. Aquatic insects, 36, 1–8.

Gamboa, M., & Watanabe, K. (2019). Genome-wide signatures of local adaptation among seven stoneflies species along nationwide latitudinal gradient in japan. BMC Genomics, 20, 84.

Gattoliatt, J-L., Rutschmann, S., Monaghan, M. T., & Sartori, M. (2018). From molecular hypothesis to valid species: description of three endemic species of Baetis (Ephemeroptera: Baetidae) from the Canary Islands. Arthropod Systematics and Phylogeny, 76, 509–528.

Genung, M. A., Schweitzer, J. A., Úbeda, F., Fitzpatrick, B. M., Pregitzer, C. C., Felker-Quinn, E., & Bailey, J. K. (2011). Genetic variation and community change-selection, evolution, and feedbacks. Functional Ecology, 25, 408–419.

Gering, J. C., Crist, T. O., & Veech, J. A. (2003). Additive partitioning of species diversity across multiple spatial scales: implications for regional conservation of biodiversity. Conservation Biology, 17, 488–499.

Gill, B. A., Harrington, R. A., Kondratieff, B. C., Zamudio, K. R., Poff, N. L., & Funk, W. C. (2013). Morphological taxonomy, DNA barcoding, and species diversity in southern Rocky Mountain headwater streams. Freshwater Science, 33, 288–301.

Guindon, S., & Gascuel, O. (2003). PhyML: A simple, fast and accurate algorithm to estimate large phylogenies by maximum likelihood. Systematic Biology, 52, 696–704.

Hauer, F. R., & Lamberti, G. A. (1996). Methods in stream ecology. Elsevier Science, California, USA.

Herrera-Vasquez, J. (2008). Community structure of aquatic insects in the Esparza River, Costa Rica. Revista de Biologia Tropical, 57, 133–139.

Heino, J., Melo A. S., Siquiera, T., Soininen, J., Valanko, S., & Bini, L. M. (2015). Metacommunity organization, spatial extent and dispersal in aquatic systems: patterns, processes and prospects. Freswater Biology, 60, 845–869.

Hughes, J. M., Schmidt, D. J., & Finn, D. S. (2009). Genes in streams: using DNA to understand the movement of freshwater fauna and their riverine habitat. BioScience, 59, 573–583.

Ishida, Y., Abekura, K., & Takemon, Y. (2005). Habitat characteristics of *Rhinogobius* sp. or “Shimahiregata” in Shirokita wando. Ecology and Civil Engineering, 8, 1–14.

Jackson, J. K., Battle, J. M., White, B. P., Pilgrim, E. M., Stein E. D., Miller, P. E., & Sweeney B. W. (2013). Cryptic biodiversity in streams: a comparison of macroinvertebrate communities based on morphological and DNA barcode identifications. Freshwater Science, 33, 312–324.

Karney CFF. (2013) Algorithms for geodesics. Journal of Geodesy. 87, 43–55.

Karaus, U., Larsen, S., Guillong, H., & Tockner, K. (2013). The contribution of lateral aquatic habitats to insect diversity along river corridors in the Alps. Landscape Ecology, 28, 1755–1767.

Kembel, S. W., Cowan, P. D., Helmus, M. R., Cornwell, W. K., Morlon, H., Ackerly, D. D.,… Webb, C. O. (2010). Picante: R tools for integrating phylogenies and ecology. Bioinformatics, 26, 1463–1464.

Lake, P.S. (2003). Ecological effects of perturbation by drought in flowing waters. Freshwater Biology, 48, 1161–1172.

Lancaster, J., & Downes, B. J. (2013). Aquatic entomology. Oxford University Press, United Kingdom.

Lande, R. (1996). Statistics and partitioning of species diversity, and similarity among multiple communities. Oikos, 76, 5–13.

Larkin, M. A., Blackshields, G., Brown, N. P., Chenna, R., McGettigan, P. A., McWilliam, H.,… Higgins, D. G. (2007). ClustalW and ClustalX version 2. Bioinformatics, 23, 2947–2948.

Legendre, P., & Anderson, M. J. (1999). Distance-based redundancy analysis: testing multispecies responses in multifactorial ecological experiments. Ecological Monographs, 69, 1–24.

Leprieur, F, Albouy, C., De Bortoli, J, Cowman, P.F., Bellwood, D.R., & Mouillot, D. (2012). Quantifying phylogenetic beta diversity: distinguishing between “true” turnover of lineages and phylogenetic diversity gradients. PlosOne, 7, e42760.

Librado, P., & Rozas, J. (2009). DnaSp v5: A software for comprehensive analysis of DNA polymorphism data. Bioinformatics, 25, 1451–1452.

Lososova, Z., Smarda, P., Chytry, M., Purschke, O., Pysek, P., Sadlo, L.,… Winter, M. (2015). Phylogenetic structure of plant species pools reflects habitats age on the geological time scale. Journal of Vegetation Science, 24, 820–833.

Lu, H.-P., Wagner, H.H., & Chen, X.-Y. (2007) A contribution diversity approach to evaluate species diversity. Basic and Applied Ecology, 8, 1–12.

Marten, A., Brandle, M., & Brandl, R. (2006). Habitat type predicts genetic population differentiation in freshwater invertebrates. Molecular Ecology, 15, 2643–2651.

Mayfield, M. M., & Levine, J. M. (2010). Opposing effects of competitive exclusion on the phylogenetic structure of communities. Ecology Letters, 13, 1085–1093.

McLain, D. K., Wesson, D. M., Oliver, J. H., & Collins, F. H. (1995). Variation in ribosomal DNA internal transcribed spacers 1 among eastern populations of *Ixodes scapularis* (Acari: Ixodidae). Journal of Medical Entomology, 32, 353–360.

Macneale, K. H., Peckarsky, B. L., & Likens, G. E., (2005). Stable isotopes identify dispersal patterns of stonefly populations living along stream corridors. Freshwater Biology, 50, 1117–1130.

Misof, B., Shanlin, L., Meusemann, K., Peters, R. S., Donath, A., Mayer C.,… Zhou, X. (2014). Phylogenomics resolves the timing and pattern of insect evolution. Science, 346, 763–767.

Murria, C., Bonada, N., Arnedo, M. A., Prat, N., & Vogler, A. P. (2013). Higher β-and γ-diversity at species and genetic levels in headwaters than in mid-order streams in *Hydropsyche* (Trichoptera). Freshwater Biology, 58, 2226–2236.

Mykra, H., Heino, J., & Muotka, T. (2007). Scale-related patterns in the spatial and environmental components of stream macroinvertebrate assemblage variation. Global Ecology and Biogeography, 16, 149–159.

Mynott, J. H., Webb, J. M., & Suter, P. J. (2011). Adult and larval associations of the alpine stonefly genus *Riekoperla* McLellan (Plecoptera: Gripopterygidae) using mitochondrial DNA. Invertebrate Systematics, 25, 11–12.

Oksanen, J., Blanchet, F. G., Kindt, R., Legendre, P., Minchin, R. B., O’Hara, R. B.,… Wagner H. (2012). Vegan. Community Ecology R Package. Retrieved from http://CRAN.R-project/package-vegan

Pastuchova, Z., Lehotsky, M., & Greeskova, A. (2008). Influence of morphohydraulic habitat structure on invertebrate communities (Ephemeroptera, Plecoptera and Trichoptera). Biologia, 63, 720–729.

Peres-Neto, P.R., Legendre, P., Dray, S. & Borcard, D. (2006). Variation partitioning of species data matrices: estimation and comparison of fractions. Ecology, 87, 2614–2625.

Prenda, J. & Gallardo-Mayenco, A. (1999). Distribution patterns, species assemblages and habitat selection of the stoneflies (Plecoptera) from two Mediterranean river basins in South Spain. International Review of Hydrobiology, 84, 595–608.

Pons, J., Barraclough, T. G., Gomez-Zurita, J., Cardoso, A., Duran, D. P., Hazell S.,… Vogler, A. P. (2006). Sequence-based species delimitation for the DNA taxonomy of undescribed insects. Systematic Biology, 55, 595–609.

Posada, D. (2008). jModelTest: Phylogenetic Model Averaging. Molecular Biology and Evolution, 25, 1253–1256.

R Core Team (2014). R: A language and environment for statistical computing. R Foundation for Statistical Computing, Vienna, Austria. URL http://www.R-project.org/.

Revell, L. J. (2012). Phytools: An R package for phylogenetic comparative biology (and other things). Methods in Ecology and Evolution, 3, 217–223.

Ribera, I., & Vogler, A. P. (2008). Habitat type as a determinant of species range sizes: the example of lotic-lentic differences in aquatic Coleoptera. Biological Journal of the Linnean Society, 71, 33–52.

Rolls, R. J., Leigh, C., & Sheldon, F. (2012). Mechanistic effects of low-flow hydrology on riverine ecosystems: ecological principles and consequences of alteration. Freshwater Science, 31, 1163–1186.

Rutschmann S., Detering, H., Simon S., Funk, D. H., Gattolliat J.-L., Hughes, S. J.,… Monaghan, M. T. (2016). Colonization and diversification of aquatic insects on three Macaronesian archipelagos using 59 nuclear loci derived from a draft genome. Molecular Phylogenetics and Evolution, 107, 27–38.

Sargent, R. D., & Ackerly, D. D. (2008). Plant-pollinator interactions and the assembly of plant communities. Trends in Ecology and Evolution, 23, 123–130.

Saito, V. S., Soininen, J., Fonseca-Gessner, A. A., & Siquiera, T. (2015a). Dispersal traits drive the phylogenetic distance decay of similarity in neotropical stream metacommunities. Journal of Biogeography, 42, 2101–2111.

Saito, V. S., Siquiera, T., & Fonseca-Gessner, A. A. (2015b). Should phylogenetic and functional diversity metrics compose macroinvertebrate multimetric indices for stream biomonitoring? Hydrobiologia, 745, 167–179.

Saito, V. S., Cianciaruso, M. V., Siqueira, T., Fonseca-Gessner, A. A., & Povoine, S. (2016). Phylogenies and traits provide distinct insights about the historical and contemporary assembly of aquatic insects communities. Ecology and Evolution, 6, 2925–2937.

Serrana, J., Miyake, Y., Gamboa, M., & Watanabe, K. (2019). Comparison of DNA metabarcoding and morphological identification for stream macroinvertebrate biodiversity assessment and monitoring. Ecological Indicators, 101, 963–972.

Sobral, F. L., & Cianciaruso, M. V. (2016). Functional and phylogenetic structure of forest and savanna bird assemblages across spatial scales. Ecography, 39, 533–541.

Stewart, K. W., & Stark, B. P. (2008). Plecoptera. In An introduction to the aquatic insects of North America (Eds. Merritt, R. W., Cummins, K. W., & Berg, M. B.), pp. 311–384. Iowa: Kendall/Hunt Publishing Co.

Thompson, R., & Townsend, C. (2006). A truce with neutral theory: local deterministic factors, species traits and dispersal limitation together determine patterns of diversity in stream invertebrates. Journal of Animal Ecology, 75, 476–484.

Tiziano, B., Fenoglio, S., Lopez-Rodriguez, M. J., Tierno de Figueroa, J. M., Grenna, M., & Cucco, M. (2010). Do predators condition the distribution of prey within micro habitats? An experiment with stoneflies (Plecoptera). Journal Review of Hydrobiology, 95, 285–295.

Tockner, K., & Stanford, J. A. (2002). Riverine flood plains: present state and future trends. Environmental Conservation, 29, 308–330.

Tockner, K., Ward, J. V., Arscott, D. B., Edwards, P. J., Kollmann, J., Gurnell, A. M.,… Maiolini, B. (2013). The Tagliamento River: A model ecosystem of European importance. Aquatic Science, 65, 239–253.

de Vienne, D. M., Giraud, T., & Martin, O. C. (2007). A Congruence Index for Testing Topological Similarity between Trees. Bioinformatics, 23, 3119–3124.

Violle C., Nemergut, D. R., Pu, Z., & Jiang, L. (2011). Phylogenetic limiting similarity and competitive exclusion. Ecology Letters, 14, 782–787.

Viteck, S., Vincon, G., Graf, W., & Pauls, S. U. (2017). High cryptic diversity in aquatic insects: an integrative approach to study the enigmatic *Leuctra inermis* species group (Plecoptera). Arthropod Systematics & Phylogeny, 75, 497–521.

Ward, J. V., Tockner, K., Edwards, P. J., Kollmann, J., Bretschko, G., Gurnell, A. M.,… Rossaro, B. (1999). A reference system for the Alps: the ‘Fiume Tagliamento’. River Research and Applications, 15, 63–75.

Webb, C. O., Ackerly, D. D., McPeek, M. A., & Donoghue, M. J. (2002). Phylogenies and community ecology. Annual Review of Ecology and Systematics, 33, 475–505.

Wiens, J. J., Ackerly, D. D., Allen, A. P., Anacker, B. L., Buckley, L. B., Cornell, H. V.,… Stephens, P. R. (2010). Niche conservatism as an emerging principle in ecology and conservation biology. Ecology Letters, 13, 1310–1324.

Xiu, J., Chen, Y., Zhang, L., Chai, Y., Wang, M., Guo Y.,… Yue, M. (2017). Using phylogeny and functional traits for assessing community assembly along environmental gradients: a deterministic process driven by elevation. Ecology and Evolution, 7, 5056–5069.

Young, M. K., Smith, R. J., Pilgrim, K. L., Fairchlid, M. P., & Schwartz, M. K. (2019). Integrative taxonomy refutes a species hypothesis: The asymmetric hybrid origin of *Arsapnia arapahoe* (Plecoptera, Capniidae). Ecology and Evolution, 9, 1364–1377.

